# ATGL is differentially required for adipocyte FABP4 secretion *in vivo* and *ex vivo*

**DOI:** 10.1101/2022.10.14.512277

**Authors:** Kacey J. Prentice, Alexandra Lee, Paulina Cedillo, Karen E. Inouye, Meric Erikci Ertunc, Grace Yankun Lee, Gökhan S. Hotamışlıgil

## Abstract

Fatty acid binding protein 4 (FABP4) is linked with the pathogenesis of metabolic diseases, including diabetes and cardiovascular disease in both mice and humans. It has also been demonstrated that the levels of hormonal FABP4 are strongly associated with obesity, and secretion is stimulated under conditions of fasting and lipolysis both *in vivo* and *in vitro*. Here, we utilized adipocyte-specific deficiency of adipose triglyceride lipase (ATGL) in a mouse model (ATGL^AdpKO^) to evaluate the regulation of FABP4 secretion by lipolytic signals in the absence of actual lipolysis *in vivo*. Previously, lipolysis-induced FABP4 secretion was found to be significantly reduced upon pharmacological inhibition of ATGL, and from adipose tissue explants from ATGL^AdpKO^ mice. Unexpectedly, upon activation beta-adrenergic receptors, ATGL^AdpKO^ mice exhibited significantly higher levels of circulating FABP4 as compared to ATGL^fl/fl^ controls *in vivo*, with no corresponding increase in non-esterified free fatty acids or glycerol, confirming the lack of lipolysis. We also generated an additional model with adipocyte-specific deletion of FABP4 in the background of ATGL^AdpKO^ mice (ATGL/FABP4^AdpKO^ or DKO) to evaluate the cellular source of circulating FABP4. In these animals, there was no evidence of lipolysis-induced FABP4 secretion, indicating that the elevated FABP4 hormone levels in the ATGL^AdpKO^ mice were indeed from the adipocytes. ATGL^AdpKO^ mice did not exhibit an increase in insulin secretion upon stimulation of lipolysis, but had a normal insulin response to glucose injection along with increased FABP4 secretion, suggesting the elevated FABP4 secretion is not due to lack of insulin. Inhibition of sympathetic signaling during lipolysis using hexamethonium significantly reduced FABP4 secretion in ATGL^AdpKO^ mice compared to controls. Therefore, activity of a key enzymatic step of lipolysis mediated by ATGL, *per se*, is not required for stimulated *in vivo* FABP4 secretion from adipocytes, which can be induced through sympathetic signaling.

## Introduction

FABP4 is a recently identified adipokine, and the only known adipokine to be secreted in response to lipolysis, with its circulating levels being increased during fasting and suppressed by insulin (1,2). Hormonal FABP4 has a strong positive correlation with BMI, and is associated with numerous metabolic disorders including diabetes, cardiovascular disease, and various types of cancers (3). Studies investigating the mechanism of action of hormonal FABP4 are yet to identify a defined receptor, however, they clearly demonstrate roles in numerous tissue types, including the liver to regulate hepatic glucose production (HGP) and the beta cell to influence insulin secretion (1,4,5). Thus, the coupling of FABP4 secretion with nutrient status likely indicates a role for FABP4 in the regulation of inter-organ communication to coordinate energy and substrate fluxes and maintain metabolic health. However, in the context of obesity where lipolysis is unrestrained due to insulin resistance, levels of FABP4 are chronically elevated in the circulation (6–8). This aberrant FABP4 secretion stimulates HGP, impairs beta cell function and potentiates the metabolic abnormalities associated with obesity (9). This role for hormonal FABP4 has been supported by studies using anti-FABP4 antibodies, which have demonstrated great therapeutic benefit in the treatment of both Type 1 and Type 2 diabetes in preclinical models (5,10,11).

Various groups have investigated the mechanisms of FABP4 secretion from adipocytes. We and others have demonstrated that FABP4 secretion is coupled to lipolytic stimuli, downstream of cAMP generation and Ca^2+^ influx (2,12,13). Consistent with this, genetic or pharmacological inhibition of adipocyte triglyceride lipase (ATGL), the first lipase involved in the lipolysis response that is responsible for the conversion of triglycerides into diglycerides (14), abolishes FABP4 secretion from adipocyte cell lines. Inhibition of hormone sensitive lipase (HSL), the second enzyme in the pathway, blunts FABP4 secretion to a lesser degree, while deletion of the final enzyme, monoacylglyceride lipase (MGL), has no effect on secretion, suggesting that initiation of lipolysis is essential for FABP4 secretion (2). Many have hypothesized that this indicates that FABP4 secretion is initiated by binding to the fatty acids liberated during this process. However, the impact of lipid exposure to increase FABP4 secretion is rather limited compared to the effect of lipolytic signals (2), suggesting the potential existence of additional regulatory mechanisms. Once lipolysis signals are delivered, FABP4 is released from adipocytes in a non-classical manner through secretory lysosomes, as evidenced by inhibition of secretion by the lysosomal secretory inhibitor, chloroquine (15).

Studies on the mechanisms of FABP4 secretion *in vivo* are very limited. In this study we aimed to investigate the role of lipolysis in FABP4 secretion *in vivo* in a model of ATGL deficiency in adipocytes. Since ATGL is the most proximal catalytic step of the lipolytic process, its deficiency in adipocytes provides a system to investigate the requirement for the catalytic process in lipolysis-induced FABP4 secretion from these cells. Our results indicate that ATGL-mediated lipid breakdown in adipocytes is not necessary for FABP4 release into circulation *in vivo*, which can be triggered through sympathetic signaling.

## Materials and Methods

### Animal Care

Animal care and experimental procedures were performed with approval from the Harvard Medical School Standing Committee on Animals. Mice were studied at 8-10 weeks of age and were maintained on a 12-hour-light/12-hour-dark cycle in the Harvard T.H. Chan School of Public Health pathogen-free barrier facility with free access to water and to a standard laboratory chow diet (PicoLab Mouse Diet 20 #5058, LabDiet). ATGL^fl/fl^ mice on a C57BL/6 background (kind gift from Dr. Erin Kershaw, University of Pittsburgh, Pittsburgh, PA) were crossed with Adiponectin-Cre mice (B6.FVB-Tg(Adipoq-cre)1Evdr/J, Jackson Labs stock #028020) to generate adipocyte-specific ATGL knockout mice (ATGL^AdpKO^) and littermate floxed controls. Generation of adipocyte-specific FABP4-ATGL-knockout mice (DKO) was accomplished by first crossing humanized FABP4 flox mice on C57BL/6 background (generated for the Hotamışlıgil lab by GenOway) with ATGL^fl/fl^ animals (2). Homozygous FABP4^fl/fl^ATGL^fl/fl^ mice were then crossed with Adiponectin-Cre mice (B6.FVB-Tg(Adipoq-cre)1Evdr/J, Jackson Labs stock #028020) to generate DKO and FABP4^fl/fl^ATGL^fl/fl^ littermate controls. Genotyping for FABP4 flox, ATGL flox, and Cre recombinase was commercially outsourced and performed from tail biopsies or ear punches using real time PCR (Transnetyx, Cordova, TN). For evaluation of body composition, mice were anesthetized with isoflurane and fat and lean mass measurements were performed using a DEXA scanner (Piximus GE).

### Adipose Explant Lipolysis

Perigonadal adipose tissue was collected from 6hr fasted 8-10 week-old male mice and washed with ice cold PBS. Explants were minced with scissors to obtain pieces of 2-4mm^3^. The pieces were washed 5 times with PBS and 5 pieces were manually picked into each well of a 12-well plate. One ml of fresh serum-free DMEM was added to each well and incubated for 1hr at 37°C. Conditioned media was harvested for the assessment of basal/unstimulated secretion. Pieces were rinsed once with fresh DMEM and DMEM containing forskolin (fsk; 1-3 μM), isoproterenol (iso; 10 μM), or norepinephrine (NE; 1μM) were added to stimulate lipolysis. The plates were incubated for 1h at 37°C, after which conditioned media was collected for measurement of FABP4 and glycerol. Adipose tissue pieces in each well were then lysed for normalization of secretion to total protein. Conditioned media was centrifuged at 5,000rpm for 10min and supernatant was used for subsequent assays.

### *In vivo* lipolysis

Lipolysis was induced in 8-10 week-old mice using the pan-beta adrenergic agonist, isoproterenol, or the beta 3-specific agonist, CL-316,243. Mice were fasted beginning at 9am for 6 hours, and lipolysis was induced by injection of freshly prepared isoproterenol (10mg/kg, IP, Tocris Bioscience, cat# 1747) or CL-316,243 (0.1mg/kg, IP, Tocris Bioscience, cat#1499) in PBS. 70μL of blood was collected from a tail nick into heparinized capillary tubes at baseline (0) before injection, and 15, 30, and 60 minutes after injection, for measurement of plasma FABP4, insulin, NEFA, and glycerol levels. For studies using hexamethonium, mice were injected IP with 20mg/kg hexamethonium (Tocris, cat#4111) or PBS 30 min prior to induction of lipolysis with CL-316,243.

### Cell Culture

3T3-L1 preadipocytes (Zenbio SP-L1-F) were maintained in DMEM with 10% bovine calf serum. For differentiation, cells were seeded in 12-well plates and grown to confluency. Adipocyte differentiation was induced in DMEM with 10% Fatal Bovine serum (FBS) using 500 μM 3-isobutyl-1-methylxanthine (IBMX, Sigma I5879), 5μg/ml insulin (Sigma I9278), 1 μM dexamethasone (Sigma D4902), and 2 μM rosiglitazone (Cayman 71740 for 48h. After which the cells were fed fresh DMEM with 10% FBS and 5μg/ml insulin every 2 days. Lipolysis assays were performed on days 10-12 of induction. For experiments involving Atglistatin (Sigma, cat#SML1075), cells were pre-treated with 10μM in DMEM for 2hrs. For induction of lipolysis, adipocytes were washed three times in PBS and incubated in serum-free DMEM for 1 hour at 37°C to evaluate basal secretion. Cells were rinsed with serum-free DMEM and lipolysis was induced by addition of DMEM with 1μM fsk, 10 μM iso, or 10% serum from treated mice for 1hr at 37°C. Conditioned media was collected at the end of each time point, centrifuged at 5,000rpm for 10 minutes, and supernatant used for further assays.

### Protein extraction, SDS PAGE, and Western blotting

Mice were fasted for 6 hours before euthanasia and tissue harvesting. Tissues were snap frozen in liquid nitrogen until protein extraction was performed. Whole adipose tissue was homogenized in ice-cold RIPA buffer (Cell Signaling Technologies, cat# 9806) containing 2mM activated Na_3_VO_4_ and 1% protease inhibitor cocktail (Sigma, cat# P340). The homogenate was centrifuged at 15000rpm at 4°C for 25 minutes to pellet the cellular debris. The resulting supernatant was assayed for protein concentration by BCA assay (Thermo Fisher/Pierce, cat# 23225). Lysates were diluted with 4X Laemmli buffer containing beta-mercaptoethanol and boiled for 10 minutes at 95°C and subjected to SDS-polyacrylamide gel electrophoresis 4-20% (Criterion TGX Stain-Free Protein Gel, Bio-Rad) gels. Protein was transferred to PVDF membrane using Bio-Rad Transblot Turbo semi-dry system. Membranes were blocked for at least 1 hour in TBS-T with 5% protease-free BSA. Detection antibodies were diluted in TBS-T with 5% protease-free BSA and incubated on membranes overnight at 4°C. Proteins of interest were detected using 1:1000 dilutions of ATGL antibody (CST, cat#2138), HRP-tagged anti-FABP4 antibody (clone 351.4.5E1.H3, generated in house for Hotamışlıgil lab by Dana Farber Antibody Core), or HRP-tagged anti-β-tubulin antibody (Abcam, cat# ab21058) as a loading control. FABP4 antibody has been validated with protein lysates from FABP4-KO mice as negative control and recombinant FABP4 as positive control. HRP signal was detected with chemiluminescent substrate (Pierce SuperSignal West Pico or Femto Plus, Thermo Fisher, cat# 34580 or 34094).

### Plasma and media assays

Plasma and media FABP4 were measured by in-house FABP4 ELISA using anti-FABP4 antibodies produced for the Hotamışlıgil lab by the Dana Farber Antibody Core (clone 351.4.6H7.G3.G9 for capture, HRP-tagged clone 351.4.5E1.H3 for detection) and recombinant human FABP4 as a standard (R&D Systems, cat# DY3150-05). Plasma insulin was assessed using a Crystal Chem Ultra Sensitive Mouse Insulin ELISA, cat# 90080 according to manufacturer instructions. Plasma and media NEFA and glycerol were measured by enzymatic colorimetric assays (FujiFilm, Wako HR Series NEFA-HR(2) cat #s 999-34691, 995-34791, 991-34891, 993-35191; Sigma Free Glycerol Reagent, cat# F2468).

### Insulin and Glucose Tolerance Tests

Before intraperitoneal (IpGTT) or oral glucose tolerance tests (OGTT), mice were fasted 16hr overnight, before injection or gavage of 0.75g/kg body weight of glucose. Insulin tolerance tests (ITT) were performed following 4-hour fasting (10am-2pm). Mice were injected intraperitoneally with 0.25 IU/kg body weight of recombinant human insulin (Humulin-R, Eli Lilly, USA) prepared in sterile PBS with 0.1% BSA. For both tests, blood glucose was evaluated at 0, 15, 30, 45, 60, 90, and 120-min post-injection. For OGTTs, blood samples were collected from a tail nick into heparinized capillary tubes for evaluation of FABP4 and insulin levels.

### Statistics

Statistical analysis was performed using GraphPad Prism version 9.4.0 for MacOS, GraphPad Software, San Diego, California USA, www.graphpad.com. All data are presented as mean ± SEM.

## Results

### Impact of ATGL on FABP4 secretion *in vitro* and *in vivo*

It is well established that lipolytic stimuli robustly increases FABP4 secretion from adipocytes *in vitro*, *ex vivo*, and *in vivo* (2,12,13,15). Activation of beta-adrenergic receptors using isoproterenol (iso), or direct stimulation of cAMP generation through activation of adenylyl cyclase using forskolin (fsk), potently stimulates lipolysis and FABP4 release from epididymal adipose tissue explants (Figure 1a). Similarly, injection of isoproterenol into wildtype mice robustly elevates plasma FABP4 levels (Figure 1b). Consistent with our previously reported results, induction of FABP4 secretion from adipocytes was dependent on ATGL activity, as treatment of 3T3-L1 adipocytes (Figure 1c) or adipose explants (Figure 1d) with Atglistatin (ATGLi), an inhibitor of ATGL and the first step of triglyceride breakdown, consistently prevents lipolysis-induced FABP4 secretion *in vitro* and *ex vivo* (2).

**Figure 1.**
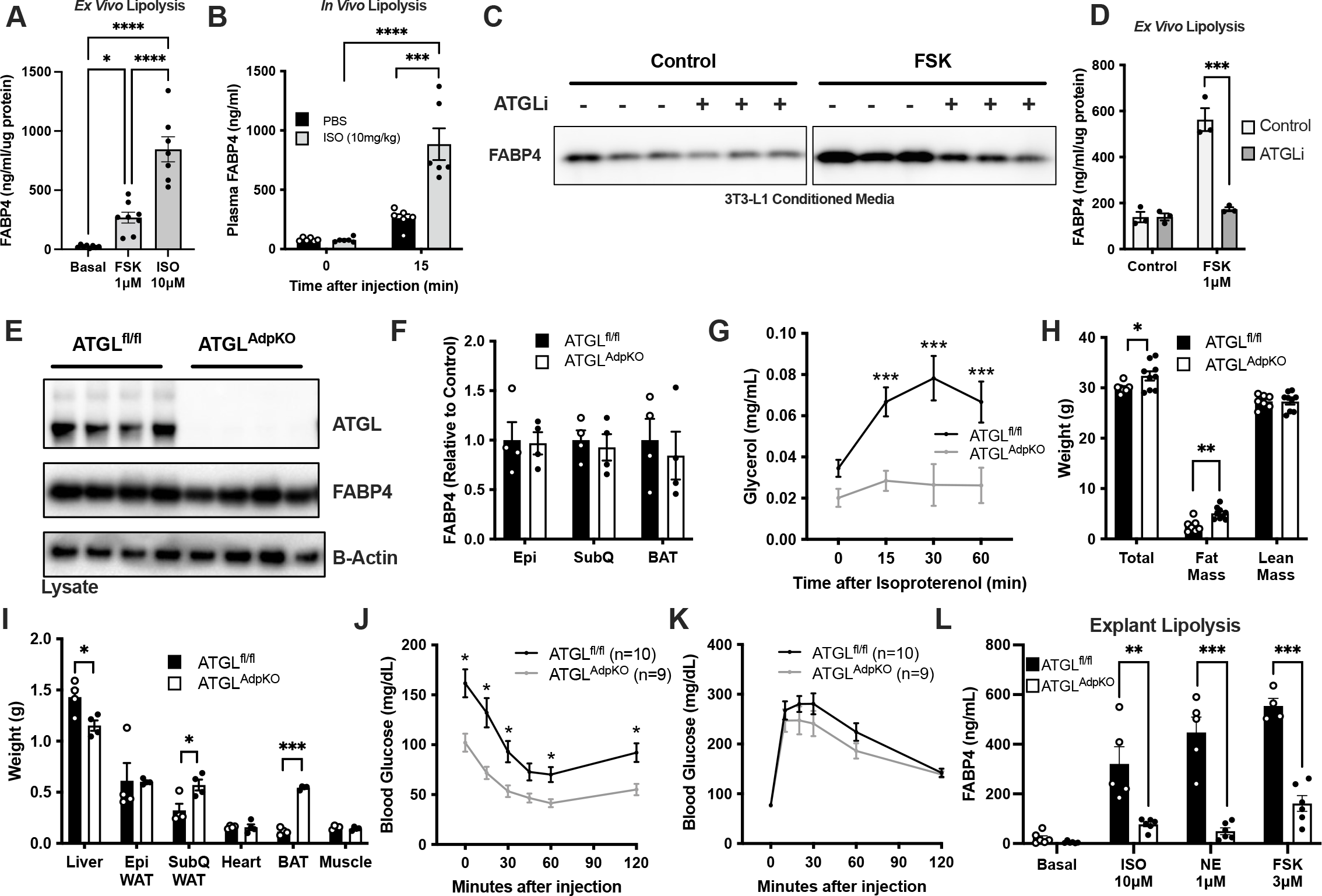
FABP4 secretion is dependent on lipolysis *in vitro*. (a) FABP4 secretion from wildtype epididymal adipose tissue explants under unstimulated (basal) conditions or induction of lipolysis by forskolin (FSK, 1 μM) or isoproterenol (ISO, 10 μM) (N=7-8/condition). (b) Plasma FABP4 in wildtype mice following injection of PBS or ISO (10mg/kg) (N=6/group). FABP4 in conditioned media from (c) 3T3-L1 adipocytes (Western blot) and (d) epidydimal adipose tissue explants (ELISA quantification) following 2hr pre-treatment with Atglistatin (ATGLi) (N=3/group). (e) Western blot of epididymal adipose tissue confirming ATGL deletion in ATGL^AdpKO^ (N=4/group). (f) Quantification of FABP4 in adipose depots from ATGL^fl/fl^ and ATGL^AdpKO^ mice (N=4/group). (g) Serum glycerol following induction of lipolysis *in vivo* (N=14-16/group). (h) Body composition as determined by DEXA (N=7-9/group). (i) Weight of liver, epididymal (Epi) adipose tissue, subcutaneous (SubQ) adipose tissue, heart, brown adipose tissue (BAT), and gastrocnemius muscle (N=4/group). Glucose excursion curves in (j) Insulin tolerance test (ITT) and (k) intraperitoneal glucose tolerance test (ipGTT) (N=9-10/group). (l) FABP4 secretion from epididymal adipose tissue explants under unstimulated (basal) conditions or induction of lipolysis by ISO (10 μM), norepinephrine (NE, 1 μM) or FSK (3 μM) (N=4-6/condition).

To gain insight into the relationship between FABP4 secretion and lipolysis *in vivo*, we generated a mouse model of adipocyte-specific ATGL-deficiency (ATGL^AdpKO^) by crossing Adiponectin-Cre expressing mice to ATGL^fl/fl^ animals. We validated the absence of ATGL protein in epididymal, subcutaneous, and brown adipose depots by western blot analysis (Figure 1e; epidydimal adipose representative). Importantly, ATGL deficiency did not impact FABP4 expression in any of the depots examined (Figure 1f). Absence of ATGL activity was confirmed by a lack of induction of lipolysis, as evaluated by plasma glycerol, following isoproterenol injection (Figure 1g). Consistent with previous reports (16,17), ATGL^AdpKO^ mice exhibited only a minor increase in body weight, attributed to increased fat mass, with no change in total lean mass (Figure 1h). Specifically, we observed increases in subcutaneous and brown fat depots, with no change in epidydimal fat mass, and a reduction in liver weight (Figure 1i). Despite a minor increase in adiposity, ATGL^AdpKO^ mice exhibited a significant improvement in insulin sensitivity, as assessed by insulin tolerance test (ITT) (Figure 1j). Furthermore, upon intraperitoneal glucose tolerance test (ipGTT), ATGL^AdpKO^ mice had a minor improvement in overall glucose tolerance (Figure 1k). Consistent with our previous observations, epididymal adipose tissue explants from ATGL^AdpKO^ mice failed to secrete FABP4 in response to lipolytic stimuli isoproterenol, fsk, or norepinephrine (NE) *ex vivo* (Figure 1l).

### Adipocyte FABP4 secretion is increased in ATGL^AdpKO^ mice *in vivo*

To evaluate if FABP4 secretion is indeed altered in the ATGL^AdpKO^ mice, consistent with *ex vivo* observations, we determined circulating FABP4 levels in response to fasting. Unexpectedly, 24hr fasted ATGL^AdpKO^ mice did not exhibit a defective induction of FABP4 secretion as compared to ATGL^fl/fl^ controls (Figure 2a). This was in the context of similar blood glucose levels and body weight following fasting (Figure 2b,c). In line with improved insulin sensitivity, ATGL^AdpKO^ mice exhibited reduced plasma insulin levels when fed *ad libitum*, however there was no difference in plasma insulin in either the fasted or re-fed states (Figure 2d). Next, we examined the lipolysis-induced FABP4 secretion in these mice, upon administration of isoproterenol to stimulate lipolysis. Strikingly, ATGL^AdpKO^ mice exhibited a marked increase in plasma FABP4 concentrations in response to lipolysis stimulation as compared to the ATGL^fl/fl^ controls, despite no induction of lipolytic activity (Figure 2e, 1g). Insulin can act to suppress lipolysis and has been shown previously to suppress FABP4 secretion *in vitro* (2). Interestingly, upon stimulation of lipolysis, ATGL^AdpKO^ mice had a significantly blunted insulin secretion response (Figure 2f), which may be associated with improved insulin sensitivity. These findings raise multiple possibilities for the mediation of hormonal FABP4 secretion in response to isoproterenol in ATGL^AdpKO^ mice *in vivo*, including the possibility of a non-adipocyte cell source, a lack of suppression by insulin, or the lack of lipolytic products.

**Figure 2.**
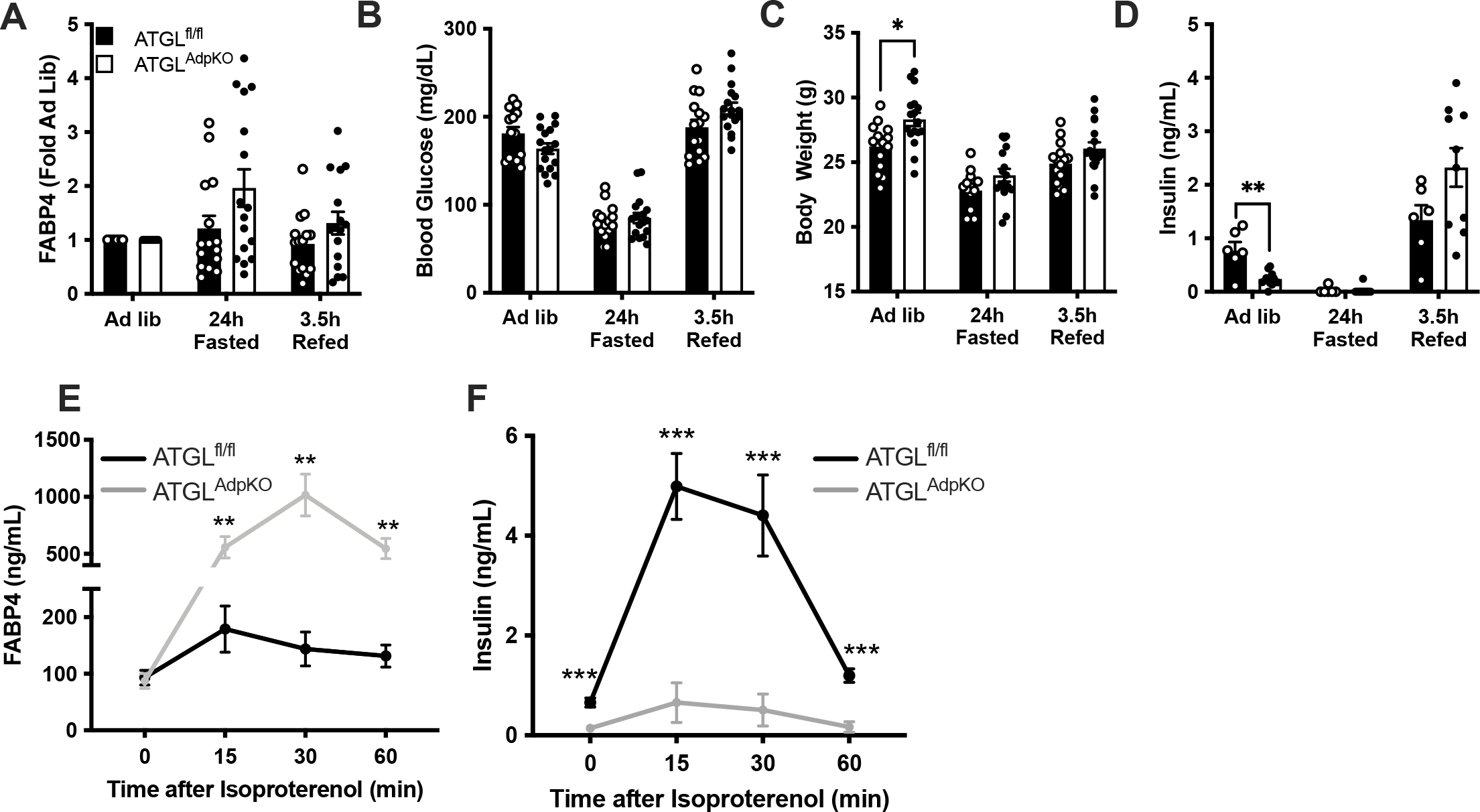
Adipocyte-specific ATGL KO mice have enhanced FABP4 secretion *in vivo*. Quantification of (a) plasma FABP4, (b) blood glucose, (c) body weight, and (d) plasma insulin in ATGL^fl/fl^ (black bars) and ATGL^AdpKO^ (white bars) mice under ad libitum fed conditions, following 24hr food withdrawl, and 3.5hrs after food was added back to cages (N=15-17/group). (e) Plasma FABP4 and (f) plasma insulin in ATGL^fl/fl^ and ATGL^AdpKO^ following induction of lipolysis with isoproterenol (10mg/kg) (N=15-16/group).

FABP4 was originally characterized as an adipocyte-specific fatty acid binding protein, however, it is now well established that it is expressed by other cell types including endothelial cells, bronchial epithelial cells, and macrophages (18–22). Given the stark contrast between *in vivo* and *ex vivo* FABP4 secretion in the ATGL^AdpKO^ mice, we investigated whether the *in vivo* lipolysis-induced FABP4 secretion may be coming from a non-adipocyte cell source in ATGL^AdpKO^ mice. Therefore, we crossed FABP4^fl/fl^ mice with the ATGL^fl/fl^ mice to generate FABP4^fl/fl^ATGL^fl/fl^ animals. These mice were then crossed with the Adiponectin-Cre line to generate FABP4/ATGL^AdpKO^ mice, hereafter referred to as double knockout (DKO), specifically lacking both FABP4 and ATGL in adipocytes. Western blot analysis of epididymal, subcutaneous, and brown adipose depots again confirmed the deletion of ATGL, and a significant reduction in FABP4 expression (Figure 3a,b). The residual FABP4 expression is likely due to non-adipocyte cell types present in explants, including endothelial cells. As observed in the ATGL^AdpKO^ mice, the DKO animals exhibited a blunted lipolysis response, confirming deletion of ATGL (Figure 3c). Similar to ATGL^AdpKO^ mice, DKO animals also exhibited a significant increase in fat mass, despite no differences in total body weight and lean mass (Figure 3d). This increase in fat mass corresponded to significant increases in both epidydimal and subcutaneous adipose depots, as well as increased brown adipose mass, with a reduction in liver weight (Figure 3e). As observed in ATGL^AdpKO^, these mice exhibited a significant improvement in insulin sensitivity by ITT (Figure 3f). When we stimulated lipolysis in the DKO animals by isoproterenol injection, there was similar induction of FABP4 in DKO mice and controls, confirming that the source of elevated FABP4 secretion in ATGL^AdpKO^ mice is indeed from adipocytes (Figure 3g). Consistent with both ATGL^AdpKO^ mice, and our previous findings in whole-body FABP4 KO mice (4), there was blunted induction of insulin secretion in the DKO animals in response to lipolysis (Figure 3h), which likely reflects the absence of lipolysis.

**Figure 3.**
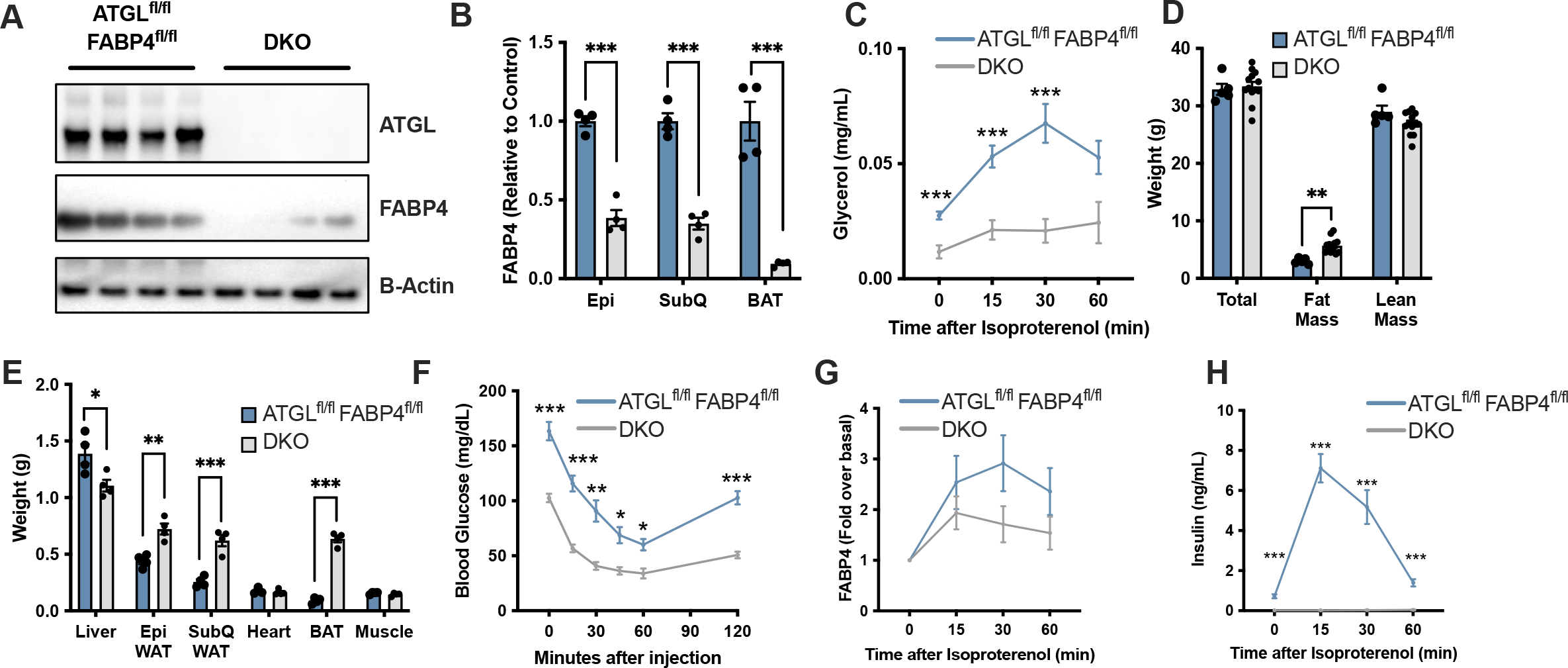
Lipolysis-induced FABP4 secretion is from adipocytes *in vivo*. (a) Western blot of epididymal adipose tissue confirming ATGL and FABP4 deletion in DKO mice (N=4/group). (b) Quantification of FABP4 in adipose depots from ATGL^fl/fl^ FABP4^fl/fl^ and DKO mice (N=4/group). (c) Serum glycerol following induction of lipolysis *in vivo* (N=13-15/group). (d) Body composition as determined by DEXA (N=5-13/group). (e) Weight of liver, epididymal (Epi) adipose tissue, subcutaneous (SubQ) adipose tissue, heart, brown adipose tissue (BAT), and gastrocnemius muscle (N=4/group). (f) Glucose excursion curves in insulin tolerance test (ITT) (N=6-9). (g) Plasma FABP4 and (h) plasma insulin in ATGL^fl/fl^ FABP4^fl/fl^ and DKO mice following induction of lipolysis with isoproterenol (10mg/kg) (N=14-15/group).

Insulin acts to suppress lipolysis through a mechanism involving reduction of intracellular cAMP, the downstream messenger activated by isoproterenol or forskolin treatment (23,24). Potentiation of adipocyte FABP4 secretion in response to lipolysis *in vivo* could be due to the lack of isoproterenol-induced insulin secretion in the ATGL^AdpKO^ mice. While insulin secretion was blunted in response to isoproterenol in the ATGL^AdpKO^ mice, we observed a normal potentiation of insulin secretion in response to glucose during an oral glucose tolerance test (OGTT), with no difference in glucose tolerance (Figure 4a,b). When we quantified plasma FABP4 10 minutes after glucose gavage we observed a significant potentiation of FABP4 secretion in both ATGL^fl/fl^ and ATGL^AdpKO^ mice (Figure 4c). Further, despite no difference in insulin secretion during OGTT, ATGL^AdpKO^ mice again secreted significantly more FABP4 than ATGL^fl/fl^ controls. Therefore, enhanced adipocyte FABP4 secretion in ATGL^AdpKO^ mice cannot be explained by a lack of insulin secretion or action.

**Figure 4.**
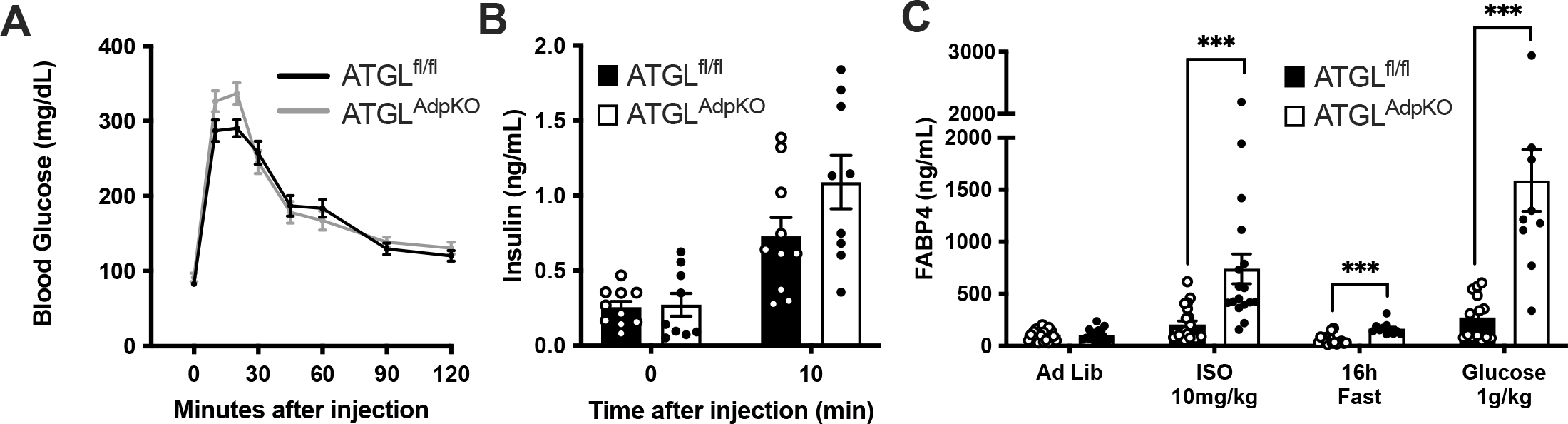
Glucose bolus stimulates FABP4 secretion *in vivo*. (a) Glucose excursion curve and (b) corresponding serum insulin levels during oral glucose tolerance test (OGTT) (N=9-10/group). (c) Plasma FABP4 in ATGL^fl/fl^ and ATGL^AdpKO^ mice under ad libidum fed, 10 minutes following isoproterenol injection (10mg/kg), 16hr fasting, and 10 minutes following glucose injection (1g/kg) (N=10-21/group).

### FABP4 secretion is potentiated by high sympathetic tone

We next speculated that there may be a circulating factor induced by isoproterenol injection *in vivo* in ATGL^AdpKO^ mice, acting to potentiate adipocyte FABP4 release, that is absent in the *ex vivo* condition. To test this, we injected ATGL^AdpKO^ mice and controls with PBS or isoproterenol and collected total serum via cardiac puncture 10 minutes later (illustrated in Figure 5a). We again validated the presence of increased isoproterenol-stimulated FABP4 levels in ATGL^AdpKO^ animals (Figure 5b). Media containing 10% serum from these mice was then added to differentiated 3T3-L1 cells and FABP4 secretion was evaluated. Serum from PBS-injected mice did not potentiate FABP4 release from adipocytes while serum from isoproterenol-injected mice did increase FABP4 release, which is likely associated with the isoproterenol contained in the serum following injection (Figure 5c). Importantly, however, there was no difference in the levels of FABP4 secretion between 3T3-L1 cells treated with serum from ATGL^fl/fl^ or ATGL^AdpKO^ mice. These data suggest that there may not be a clear circulating factor responsible for increased FABP4 release in the absence of adipocyte ATGL.

**Figure 5.**
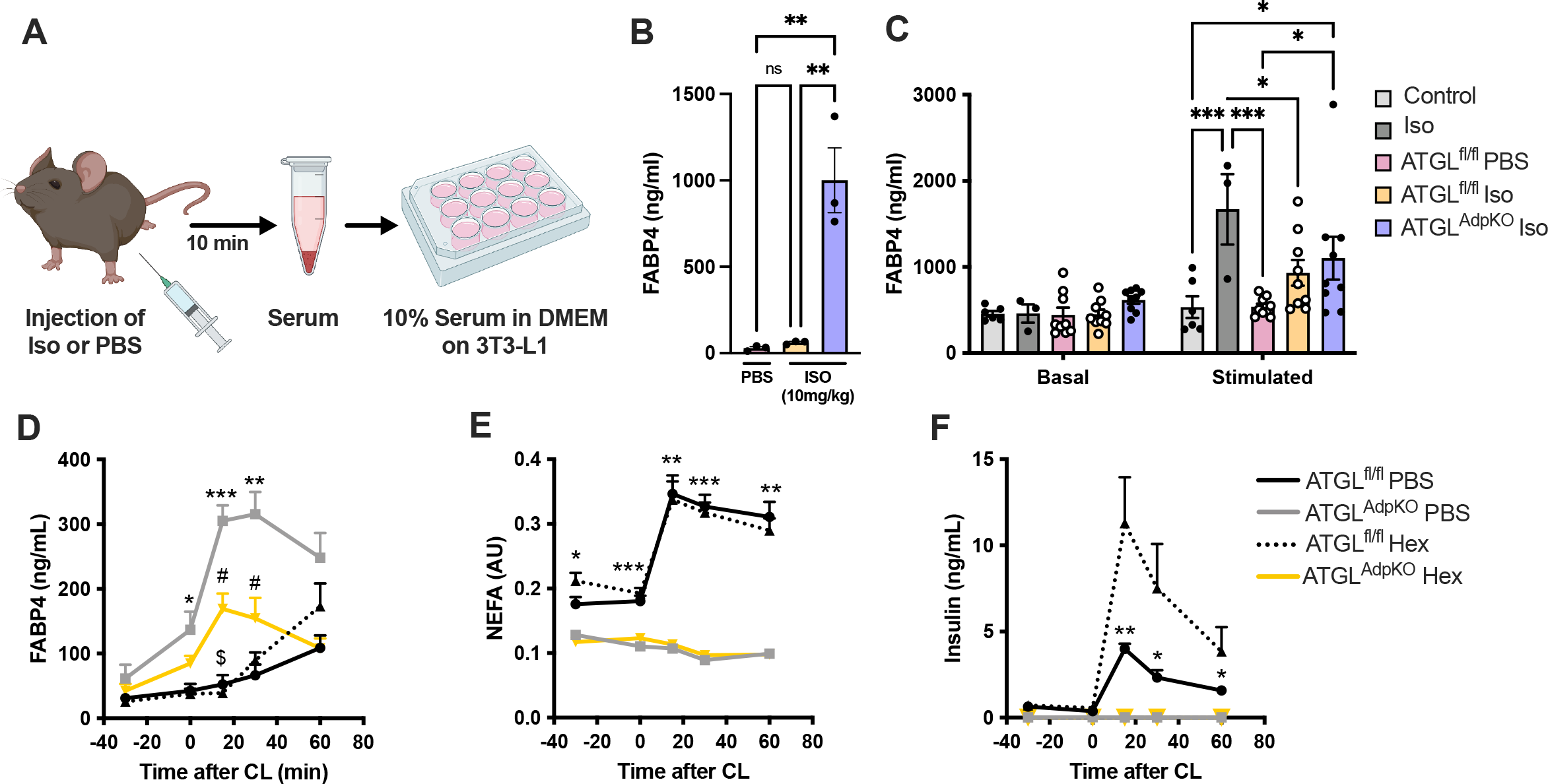
*In vivo* FABP4 secretion is blunted upon sympathetic nervous inhibition. (a) Schematic illustration of the protocol for serum induction of lipolysis in 3T3-L1 cells. (b) Serum FABP4 levels in mice 10 minutes following injection of PBS or isoproterenol (ISO, 10mg/kg) (N=3/group). (c) Quantification of FABP4 in conditioned media from 3T3-L1 cells treated with control, isoproterenol (1 μM), or 10% serum from mice in (b) (N=3-9/group). Serum (d) FABP4, (e) non-esterified fatty acids (NEFA), and (f) insulin following administration of 20mg/kg hexamethonium and 0.1mg/kg CL-316,243 (N=5-6/group).

Finally, we hypothesized that the *in vivo* signal regulating FABP4 release from adipocytes upon isoproterenol injection that is absent from *ex vivo* conditions may be related to sympathetic innervation of the tissue. Previously, we have demonstrated that FABP4 secretion upon exposure to propionic acid, *in vivo*, is diminished when sympathetic signaling is pharmacologically inhibited (25). To test if this is indeed the factor primarily regulating FABP4 release in the ATGL^AdpKO^ mice, we pre-treated mice with hexamethonium (Hex), a non-depolarizing ganglionic blocker (26), 30 minutes prior to CL-316,243 injection. Indeed, FABP4 secretion was reduced by more than 50% upon Hex administration in ATGL^AdpKO^ mice compared to ATGL^AdpKO^ mice without Hex (Figure 5d). Importantly, Hex treatment had no effect on the induction of lipolysis in ATGL^fl/fl^ mice (Figure 5e). Insulin secretion was also not altered with Hex treatment in the ATGL^AdpKO^ mice, but was significantly elevated in ATGL^fl/fl^ mice (Figure 6f). Therefore, the enhanced FABP4 secretion from ATGL^AdpKO^ mice upon lipolysis *in vivo* is likely due elevated sympathetic tone (27) resulting in altered adipocyte signaling and potentiation in FABP4 release which is absent *ex vivo*.

## Discussion

FABP4 is secreted from adipocytes upon exposure to lipolytic stimuli. Under conditions of obesity, loss of insulin sensitivity by adipocytes results in unsuppressed lipolysis, which leads to increased levels of hormonal FABP4. Circulating FABP4 levels exhibit strong positive correlation with diabetes and cardiovascular disease, and multiple independent GWAS studies have identified protection in individuals with a low expression variant of FABP4 against Type 2 diabetes and cardiovascular diseases (28–30). Importantly, targeting hormonal FABP4, either through genetic ablation or monoclonal antibody therapy has proven to protect against or reverse diabetes in preclinical models of genetic or diet-induced obesity (5,9,10). Therefore, understanding the mechanisms underlying the secretion and regulation of this hormone is of critical importance to realize its broad utility in multiple human diseases.

Previous work has provided an understanding of the *in vitro* or *ex vivo* mechanisms regulating FABP4 secretion from adipocytes. Increases in cAMP and intracellular calcium levels, downstream of beta3-adrenergic receptor activation, are essential for the potentiation of FABP4 release (2,12,13). We have previously reported that the enzymes of lipolysis, ATGL and, to a lesser extent HSL are required for FABP4 secretion *ex vivo* and *in vitro*, as inhibition of ATGL either genetically or pharmacologically abolished secretion (2). Supportive of the role for lipolytic process in the secretion of FABP4, inhibition of HSL or MGL, as the lipolytic enzymes required for the second and third steps of TG breakdown, respectively, had decreasing efficacy at reducing FABP4 secretion, suggesting that initiation of lipolysis was essential for this process. Here, however, we have shown that while lipolysis is required for *ex vivo* secretion of FABP4, there may be alternative pathways regulating secretion *in vivo*.

Our studies in the adipocyte-specific ATGL-deficient mouse have demonstrated that one such signal promoting the unexpected degree of FABP4 secretion in this setting comes from the sympathetic system, *in vivo*. Blocking sympathetic signaling resulted in significantly reduced FABP4 secretion in the context of adipocyte ATGL-deficiency, indicating that this is a critical pathway regulating secretion *in vivo*. Importantly, it is reported that adipocyte-specific ATGL knockout mice exhibit higher basal sympathetic tone than wildtype controls, which could further explain the increase we observe in FABP4 secretion in these mice (27). Similarly, human obesity is also characterized by an increase in sympathetic tone (31), which may also contribute to the observed correlation between hormonal FABP4 and BMI (21), and exacerbate the pathological elevations of circulating FABP4. The role for sympathetic activation in FABP4 release may be related to a role for this protein in so-called “fight or flight” responses, enabling the rapid release of fatty acids coupled to FABP4 to provide fuel in life threatening conditions, and to potentiate hepatic glucose production for the same purpose (3). It is also possible that the lack of availability of the products of lipolysis upon beta-adrenergic stimulation in the ATGL^AdpKO^ mice is also a source of stress in this model, potentiating the release of FABP4. The lack of lipolysis-induced insulin secretion in the ATGL^AdpKO^ model likely also contributes to the potentiation of FABP4 release, as there is an absence of counter-regulatory stimuli to the lipolysis induction. Indeed, *ex vivo* insulin acts to suppress FABP4 release from wildtype adipose tissue and adipocyte cell lines (2). However, due to the extreme insulin sensitivity of the ATGL^AdpKO^ mice, our ability to supplement insulin during conditions of lipolysis to test this question is limited.

It is also of interest that the ATGL-deficient animals remained insulin sensitive despite high levels of lipolysis-associated FABP4 in circulation. In this particular model, while the lipolytic signal is delivered to trigger FABP4 release, it occurs in the absence of lipolytic process and any associated lipid or other products. Given the strong link between hormonal FABP4 and a multitude of chronic and metabolic disease in both mice and humans, it is possible that there are different functional roles for FABP4 depending on the cellular source and the lipid environment in the biological functions of this hormone. This is an intriguing system to explore further the mechanisms and functional relevance of apo-versus holo-FABP4 *in vivo* in future studies

## Data Availability Statement

All raw data will be publicly available upon publication of the manuscript.

## Author contributions

K.J.P. designed and performed the experiments, analyzed the data, and prepared the manuscript. A.L., P.C., K.E.I., and M.E.E. assisted with designing and conducting *in vivo* experiments. A.L., P.C., and G.Y.L assisted with performing *in vitro* and *ex vivo* experiments. A.L., P.C., K.E.I., M.E.E., and G.Y.L. analyzed the data and revised the manuscript. G.S.H. supervised and supported the project, interpreted results, and revised the manuscript.

## Acknowledgements

The authors thank members of the Hotamışlıgil laboratory, past and present, for their intellectual contributions and assistance with experiments. ATGL^fl/fl^ mice were a kind gift from Dr. Erin Kershaw, University of Pittsburgh, Pittsburgh, PA.

